# Attenuation of cGAS/STING Activity During Mitosis

**DOI:** 10.1101/2019.12.19.883090

**Authors:** Brittany L. Uhlorn, Eduardo R. Gamez, Shuaizhi Li, Samuel K. Campos

## Abstract

The innate immune system recognizes cytosolic DNA associated with microbial infections and cellular stress via the cGAS/STING pathway, leading to activation of phospho-IRF3 and downstream IFN-I and senescence responses. To prevent hyperactivation, cGAS/STING is presumed to be non-responsive to chromosomal self DNA during open mitosis, though specific regulatory mechanisms are lacking. Given a role for the Golgi in STING activation, we investigated the state of the cGAS/STING pathway in interphase cells with artificially vesiculated Golgi and in cells arrested in mitosis. We find that while cGAS activity is impaired through interaction with mitotic chromosomes, Golgi integrity has little effect on the enzyme’s production of cGAMP. In contrast, STING activation in response to either foreign DNA (cGAS-dependent) or exogenous cGAMP is impaired by a vesiculated Golgi. Overall our data suggest a secondary means for cells to limit potentially harmful cGAS/STING responses during open mitosis via natural Golgi vesiculation.

## Introduction

Cells possess intrinsic sensory pathways as part of the innate immune system to detect microbial infection or other physiological insults [1]. Foreign nucleic acids are often recognized as pathogen-associated molecular patterns (PAMPs) through a number of pattern recognition receptors (PRRs) [2], causing activation of NFκB-dependent inflammatory cytokine responses and/or IRF3/7-dependent type-I interferon (IFN-I) responses [1, 3]. The cGAS/STING pathway is recognized as a central component of innate immunity for cytosolic DNA recognition and downstream IFN-I responses [4–8]. Cytosolic DNA is recognized by the enzyme cGAS, triggering production of the cyclic dinucleotide 2’,3’-cGAMP [9]. STING, a transmembrane endoplasmic reticulum (ER) protein [10, 11], is activated by direct binding to cGAMP [12].

Upon activation by cGAMP at the ER, dimeric STING undergoes a conformational change [13] and traffics to the Golgi, a prerequisite for assembly of the STING/TBK1/IRF3 complex and downstream IFN-I responses [14]. cGAMP-dependent STING recruitment of TBK1 [15] can lead to phosphorylation of IRF3 and NFκB, stimulating both IFN-I and proinflammatory cytokine responses [10, 16, 17]. Trafficking of STING to the Golgi is regulated by several host factors including iRHOM2-recruited TRAPß [18], TMED2 [19], STIM1 [20], TMEM203 [21], and Atg9a [22]. STING activation at the Golgi requires palmitoylation [23] and ubiquitylation [24, 25], allowing for assembly of oligomeric STING and recruitment of TBK1 and IRF3 [26–28]. STING also interacts with the ER adaptor SCAP at the Golgi to facilitate recruitment of IRF3 [29],

In addition to innate defense against microbial infections, cGAS/STING is involved in cellular responses to DNA damage and replicative/mitotic stress [5, 30–36]. DNA damage, replicative stress, chromosomal instability, and mitotic errors can lead to the formation of micronuclei which can trigger antiproliferative IFN-I and senescence responses via cGAS/STING [37].

Unabated activation of cGAS/STING can lead to harmful autoinflammatory and senescence responses, exemplified by type-I interferonopathies associated with mutations in STING [38–41] or mutations in the DNases TREX1 and DNASE2 that normally clear cells of cGAS-stimulatory DNA [42–45]. Given the harmful effects of cGAS/STING hyperactivation, cells need regulatory mechanisms to avoid self-stimulation of cGAS/STING during mitosis. Cytosolic compartmentalization of cGAS was initially proposed as a mechanism, but nuclear chromosomes and cytosolic compartments mix upon mitotic nuclear envelope breakdown (NEBD), suggesting a more elaborate means of cGAS/STING attentuation during cell division.

The chromatinized nature of cellular genomic DNA has been proposed to mitigate cGAS/STING activity, with histones structurally marking DNA as “self.” This is an attractive model as many DNA viruses sensed by cGAS/STING upon initial entry, trafficking, and uncoating (prior to viral DNA replication) contain either naked, unchromatinized dsDNA (e.g. herpesviridae [46, 47]) or DNA that is packaged with non-histone viral core proteins (e.g. adenoviridae, poxviridae, and asfarviridae [48–52]).

Since cGAS localizes to condensed chromosomes upon NEBD [35, 53], others have asked whether cGAS is activated by chromosomes, and if not, what mechanisms exist to prevent such self-activation. Recent studies have revealed that i) chromosome-bound cGAS is tightly tethered to chromatin, potentially via interactions with H2a/H2b dimers, ii) chromatin interaction does not involve the DNA-binding domains of cGAS required for “typical” activation by dsDNA, and iii) that chromosome binding results only in weak activation of cGAS with relatively low production of cGAMP [34, 53, 54].

Given the importance of the Golgi in STING-dependent activation of IRF3, we hypothesize a parallel mechanism for cGAS/STING regulation during open mitosis – Golgi vesiculation [55, 56]. Here, we find that chemical dispersal of the Golgi abrogates cGAS/STING-dependent phospho-IRF3 responses to transfected DNA. Furthermore, we show that cGAS/STING activity in response to transfected DNA is diminished during open mitosis, correlating with the vesiculated state of the mitotic Golgi. This Golgi-dependent weakening of cGAS/STING responses to transfected DNA occurs at the level of STING, as cGAS activity is downregulated upon mitotic chromosome binding but largely unaffected by Golgi integrity. The vesiculated state of the mitotic Golgi may therefore provide an additional safeguard mechanism, ensuring that potentially harmful cGAS/STING responses to self-DNA are minimized during cell division.

## Results & Discussion

### Human Keratinocytes Respond to Foreign DNA via cGAS/STING

We chose to investigate activity of the cGAS/STING pathway in HaCaTs, a spontaneously immortalized human keratinocyte cell line [57]. This cell line is a good model for primary keratinocytes that serve a barrier function and as host cells for a number of viral infections including papillomaviruses, herpesviruses, mosquito-transmitted alphaviruses and flaviviruses [58–64], and numerous bacterial and fungal pathogens [65, 66]. Although prior work has shown that HaCaT cells express an intact cGAS/STING pathway [60, 67–71], we sought to ensure the cGAS/STING pathway was functional and responsive in our HaCaT line, and that this pathway was the primary mode of IRF3 phosphorylation in response to foreign DNA.

HaCaTs were transfected with siRNAs targeting the cGAS/STING pathway and subsequently transfected with endotoxin-free dsDNA plasmid pGL3 (Figure 1). IRF3 was phosphorylated (pIRF3) in response to DNA transfection, exemplifying the ability of exogenous dsDNA to activate the cGAS/STING pathway in HaCaTs. IRF3 phosphorylation was impaired when pGL3 was transfected following siRNA knockdown of cGAS, STING, or IRF3, confirming that HaCaT cells utilize the cGAS/STING pathway as the primary mechanism of activating a pIRF3 response to foreign DNA.

**Figure 1.**
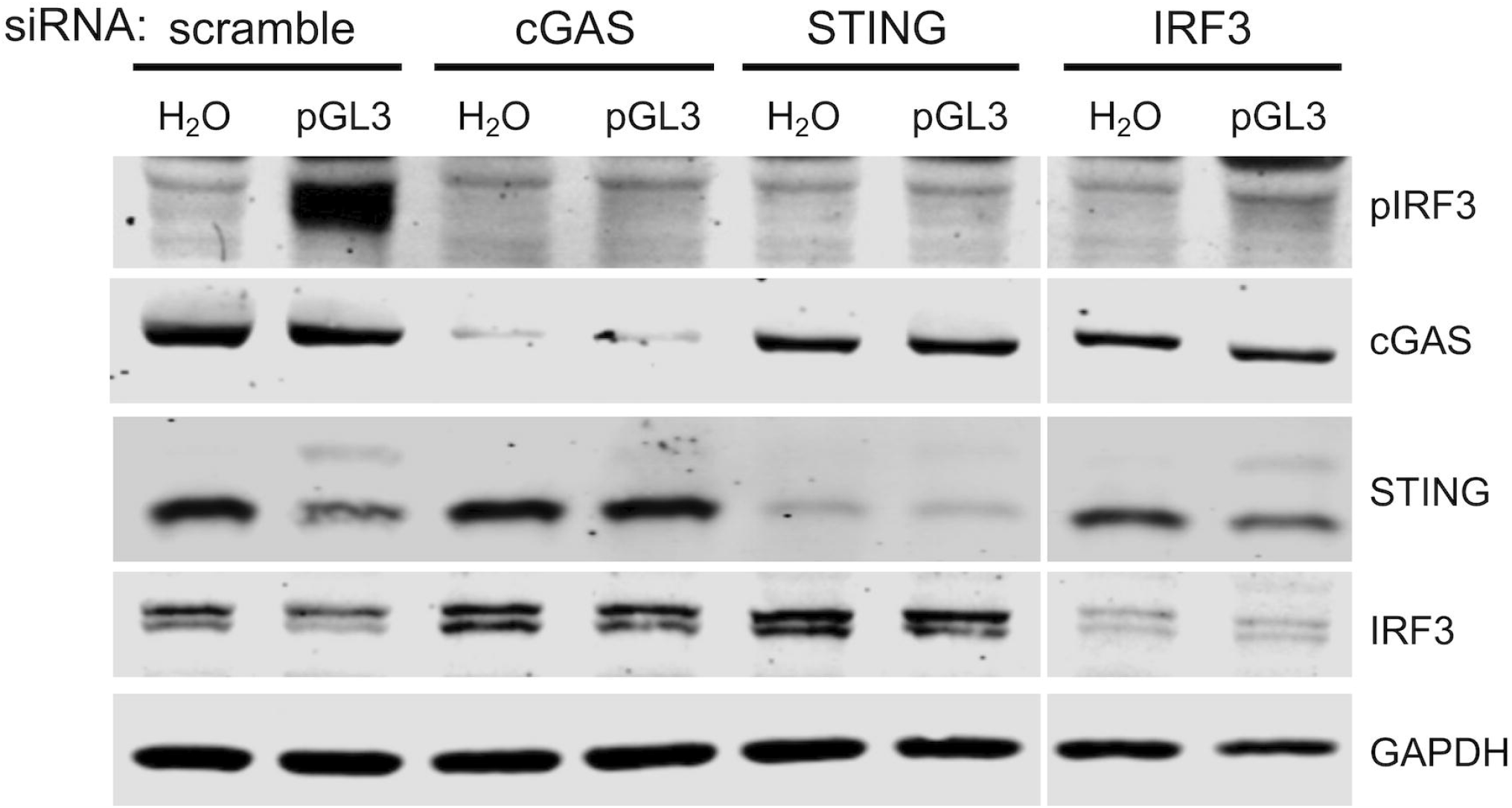
Human keratinocytes respond to foreign DNA via cGAS/STING. HaCaTs were transfected with siRNAs for 16 hr, followed by transfection with 500 ng pGL3 or water for 90 min. Lysates were analyzed for cGAS/STING component knockdown and pathway activity by SDS-PAGE and western blot.

### Golgi Vesiculation but not Fragmentation Impairs cGAS/STING Activity at the Level of STING

Activated STING traffics to the Golgi to oligomerize and complex with TBK1/IRF3. Since Golgi trafficking is critical for STING/TBK1/IRF3 complex assembly and activation, we hypothesized that a vesiculated Golgi would prevent cGAS/STING from responding to foreign DNA. To test this idea, we used the Golgi-disrupting compounds nocodazole (NOC), golgicide A (GCA), and brefeldin A (BFA). At high dose, NOC treatment causes reversible Golgi fragmentation and redistribution to ER-exit sites (ERES) [72] whereas GCA and BFA treatment cause more drastic Golgi vesiculation and dispersal by targeting the Arf1 guanine nucleotide exchange factor GBF1 [73, 74]. Indeed, NOC treatment of HaCaT cells caused a pronounced fragmentation of the Golgi as seen by immunofluorescence (IF) microscopy for trans-Golgi markers p230 and TGN46, while GCA and BFA treatment completely vesiculated the Golgi apparatus (Figure 2A, B), mimicking the dispersed mitotic Golgi. As expected, transfection of pGL3 resulted in clustering of STING with p230- and TGN46-positive Golgi membranes and nuclear import of IRF3, indicative of pathway activation (Figure 2 A-D). Addition of GCA or BFA completely abrobated activation of STING and IRF3, while NOC had minimal effect, suggesting that while a fragmented Golgi can support pGL3-stimulated STING/IRF3 activation, a dispersed Golgi cannot (Figure 2A-D). Likewise, IRF3 phosphorylation was induced upon transfection of HSV-60 oligonucleotide DNA, calf-thymus DNA (CTD), or pGL3 plasmid in HaCaTs with intact or fragmented Golgi; however, GCA- or BFA-mediated Golgi vesiculation impaired DNA-dependent IRF3 phosphorylation (Figure 3A), in agreement with prior literature [75].

**Figure 2.**
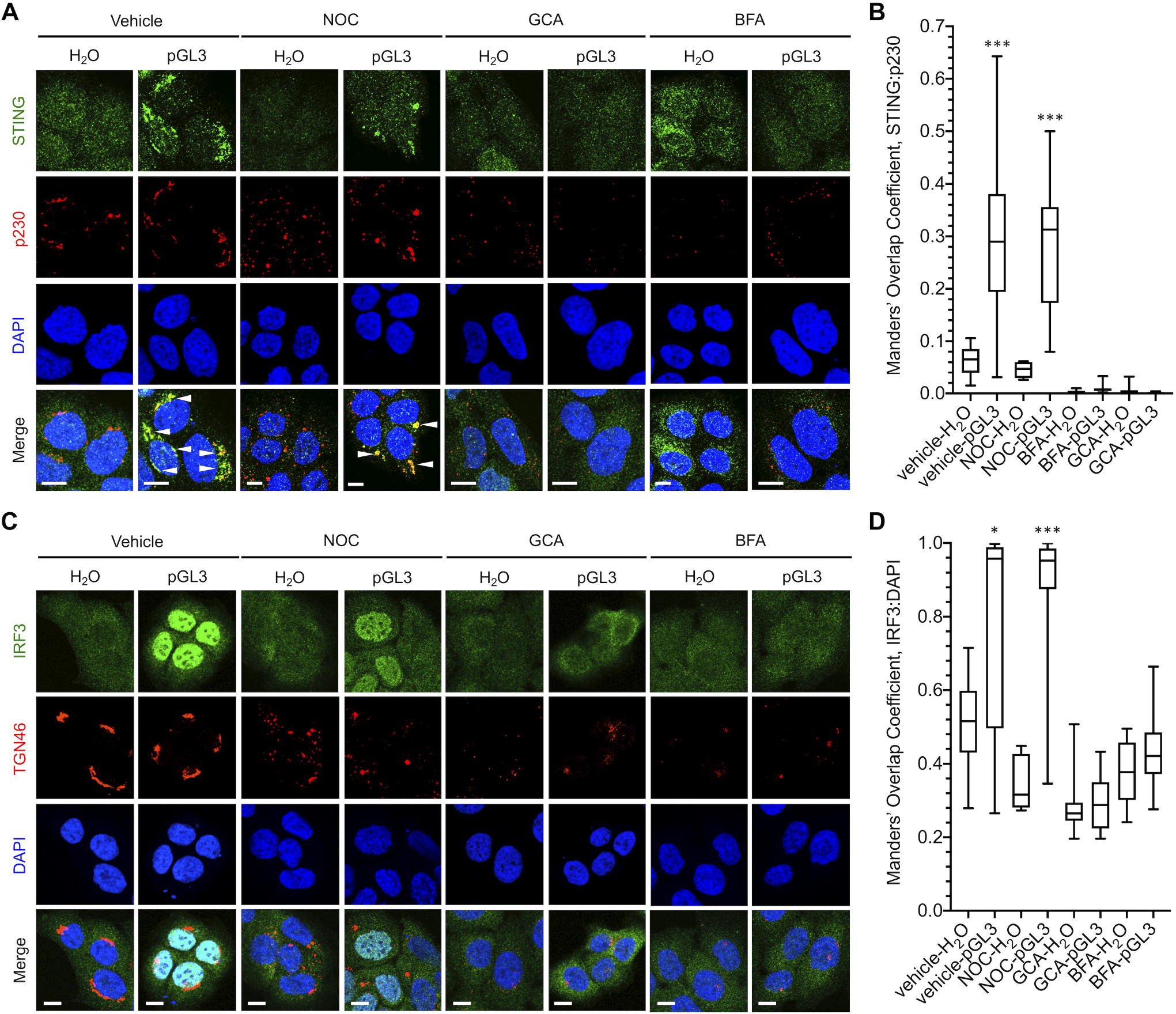
Effects of Golgi disruption on DNA-dependent subcellular localization of STING and IRF3. HaCaT cells were pretreated with vehicle, NOC, GCA, or BFA prior to a 90 min transfection with H_2_O or pGL3. Cells were fixed and stained for **(A)** STING and p230 or **(C)** IRF3 and TGN46 prior to DAPI staining. Representative micrographs are shown in **(A, C)**, white arrowheads indicate overlap. **(B, D)** Manders’ overlap coefficients from multiple micrographs were plotted for **(B)** STING:p230 and **(D)** IRF3:DAPI. *p < 0.01, ***p < 0.0001. Scale bars = 10 μm.

**Figure 3.**
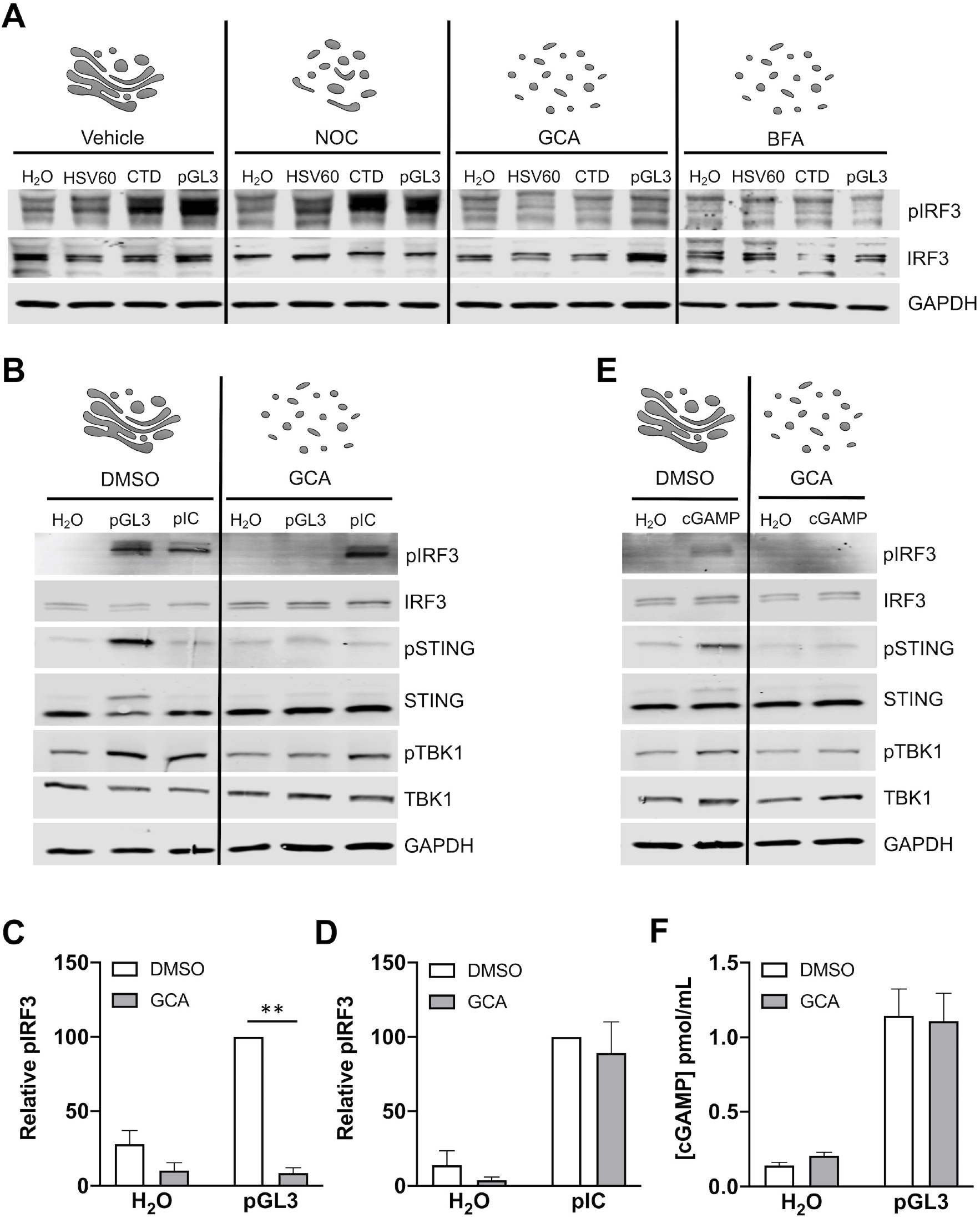
Effects of Golgi disruption on cGAS/STING activity. **(A)** Cells were treated with vehicle, nocodazole (NOC), golgicide A (GCA), or brefeldin A (BFA) for 1 hr prior and during a 90 min transfection with HSV-60 oligonucleotide, calf-thymus DNA (CTD), or plasmid pGL3. **(B, C, D)** Transfection of vehicle- or GCA-treated cells with pGL3 or pIC, and **(C, D)** densitometric quantification of pIRF3 blots. **p < 0.001, n = 5 biological replicates. **(E)** Vehicle- or GCA-treated cells were treated with H_2_O or 12.5 μg cGAMP for 90 min prior to SDS-PAGE and western blot for cGAS/STING pathway components. **(F)** cGAMP production in vehicle- and GCA--treated cells upon pGL3 transfection. n = 3 biological replicates, with n = 2 technical replicates each.

To determine if the requirement for intact Golgi was specific for cGAS/STING or involved a more broad inhibition of IRF3 phosphorylation, we investigated activation of the RIG-I-like receptors (RLRs). The dsRNA-mimic polyinosinic-polycytidylic acid (pIC) stimulates RLR-family members which recognize intracellular viral RNA products through the mitochondria-resident adaptor protein MAVS to elicit NFΚB inflammatory and IRF3/7-dependent IFN-I responses [76, 77]. pGL3-dependent pSTING, pTBK1, and pIRF3 responses were abolished by Golgi dispersal (Figure 3B, C). In contrast, transfection of pIC elicited pTBK1 and pIRF3 responses regardless of GCA treatment (Figure 3B, D), suggesting that the cGAS/STING pathway, but not the RLR pathway, is regulated by Golgi morphology. Exogenous stimulation of HaCaT cells with the STING ligand cGAMP was sensitive to GCA-mediated Golgi disruption (Figure 3E) and GCA had no effect on cGAMP production in response to pGL3 transfection (Fig. 3F), indicating that Golgi dispersal blocked the pathway downstream of cGAS.

The GCA-induced repression of cGAS/STING activation was also reflected when measuring pGL3-dependent downstream transcriptional responses. RT-qPCR at 4 hr and 8hr post pGL3 transfection revealed that induction of IFN-I (IFNB1), ISGs (Viperin, IFI6, HERC5, IFIT2, IFIT3), and chemokines (CXCL10 and CXCL11) was significantly dampened in the presence of GCA (Figure 4). Overall, these data show that cGAS/STING activity is blunted at the level of STING upon Golgi vesiculation, similar to what may occur during mitosis.

**Figure 4.**
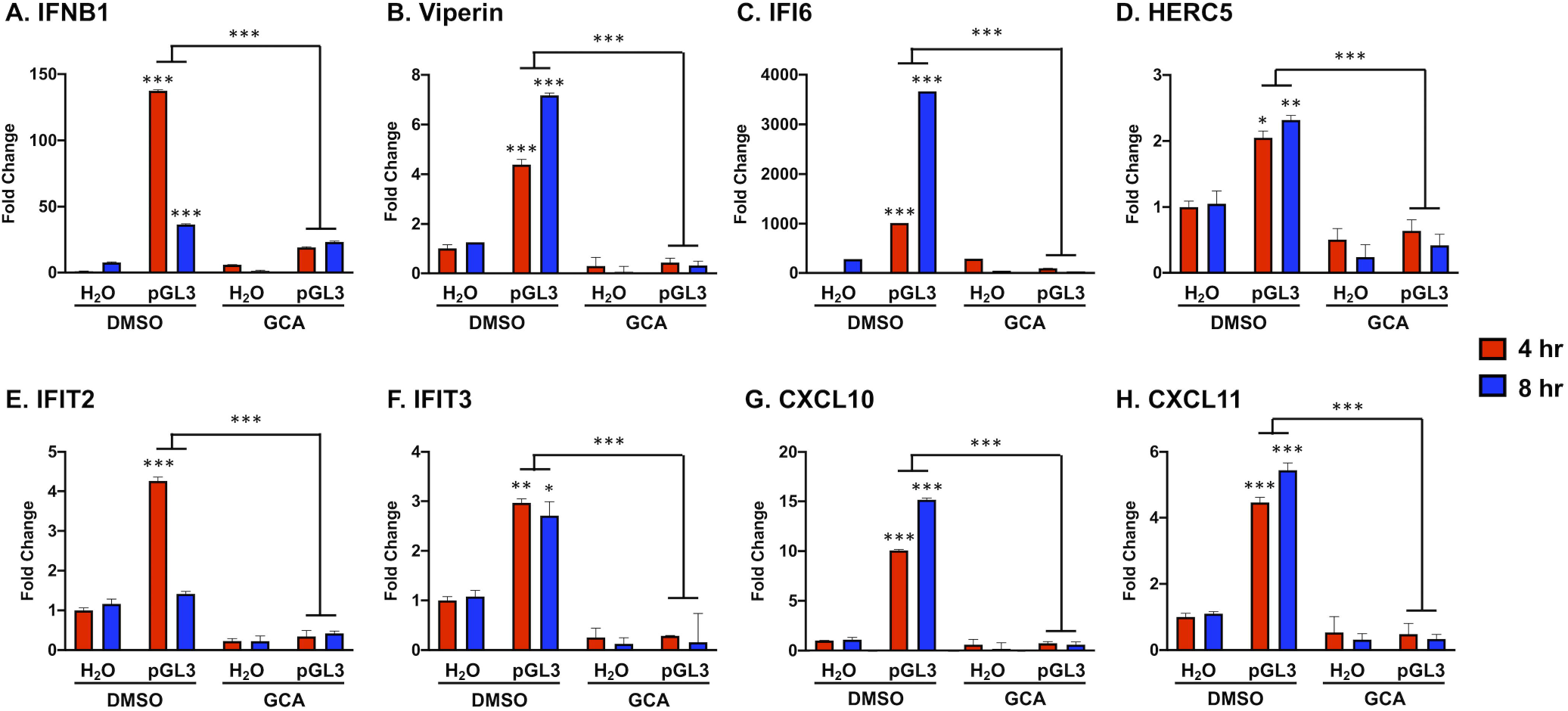
Golgi vesiculation impairs DNA-dependent IFN, ISG and chemokine gene transcription. HaCaT cells were treated with GCA or vehicle for 1 hr prior to a 90 min pGL3 transfection. Transcript levels were measured via RT-qPCR and normalized to TATA-binding protein. *p < 0.01, **p < 0.001, ***p < 0.0001, for comparisons of 4 hr and 8 hr DMSO-treated pGL3 groups to 4 hr and 8 hr DMSO-treated H_2_O controls and for 4 hr and 8 hr DMSO-treated pGL3 groups to 4 hr and 8 hr GCA-treated pGL3 groups, n = 3 technical replicates.

### cGAS/STING Activity is Attenuated During Mitosis

We next investigated the impact of natural mitotic Golgi vesiculation on the ability of cGAS/STING to sense and respond to exogenous DNA. Secretory ER to Golgi traffic is blocked during mitosis [78–81]. Golgi integrity is dependent on cargo transport from ERES, and mitotic arrest of COPII-dependent ERES traffic causes Golgi dispersal [82, 83]. As assembly of the activated STING/TBK1/IRF3 complex requires STING transport from ERES to the Golgi [14], we hypothesized that mitotic Golgi dispersal and inactivation of ERES would blunt cGAS/STING responses.

We devised a method to synchronize cells at prometaphase (Figure 5A). Briefly, cells were cultured at 100% confluence for 48 hr in 1% serum, leading to quiescent arrest in G_0_. Cells were released by replating in 10% serum, allowing for G_1_ re-entry and progression to S. Twenty-four hours post-G_0_ release, cells were synchronized at prometaphase with low dose NOC for 12 hr. Upon NOC washout, synchronized cells progressed through mitosis, returning to G_1_ within 3 hr. Propidium iodide staining showed cells enriched at G_2_/M after NOC treatment, and the majority of cells back in G_1_ by 3 hr post-NOC release (Figure 5A). Phosphorylated histone H3 (pH3) was enriched in NOC-synchronized cells, decreasing as cells returned to G_1_ (Figure 5B).

**Figure 5.**
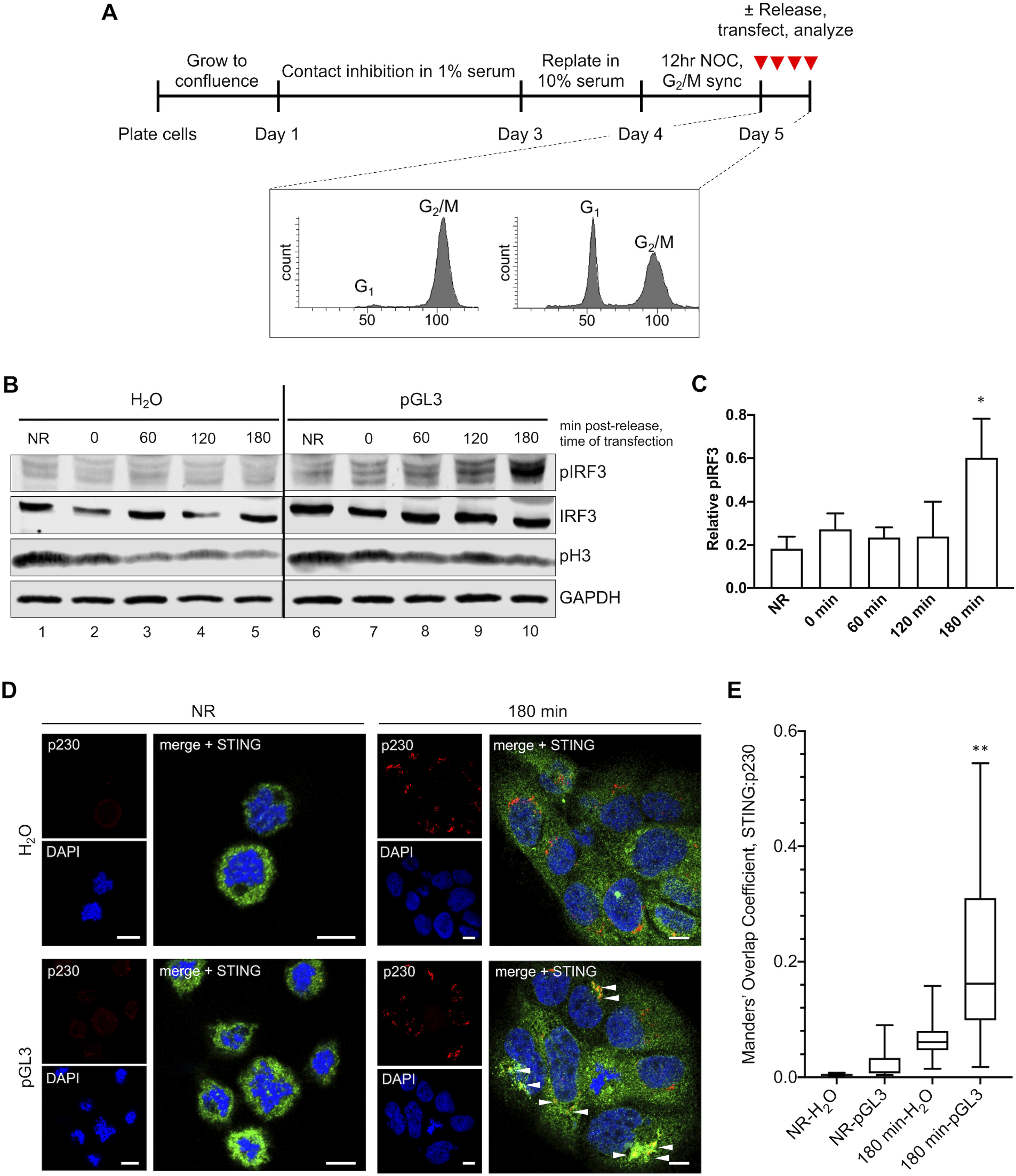
cGAS/STING activity is attenuated during mitosis. **(A)** Cell synchronization method. Cells were pre-synchronized in G_1_ by contact inhibition and growth in low serum. Cells were then released by replating at subconfluence in 10% serum and synchronized in prometaphase with NOC. Cell cycle analysis by PI-staining. Cells were transfected with pGL3 for 90 min at various times post-release from NOC. **(B)** pIRF3 activation in response to pGL3 was dampened when arrested cells were transfected, gradually returning as cells completed mitosis and returned to interphase. **(C)** Densitometric quantification of relative pIRF3 increase in response to pGL3 transfection across multiple blots. *p < 0.05, for comparison of 180 min to NR groups, n = 4 independent biological replicates. **(D)** Golgi localization of STING was impaired when arrested cells (NR) were transfected, but was restored upon return to interphase (180 min). **(E)** Manders’ overlap coefficients from multiple micrographs were plotted for STING:p230, **p < 0.001, scale bars = 10 μm.

We used this scheme to assess mitotic cGAS/STING responses to pGL3 transfection. Cells were transfected while in prometaphase following the 12 hr NOC sync (non-released group, NR), or at 0, 60, 120, or 180 min post-release, and cGAS/STING activity was assessed 90 min post-transfection. Cells arrested at prometaphase mounted a very weak pIRF3 response to pGL3 transfection compared to cells which had returned to G_1_ after a 3 hr release (Figure 5B, compare lanes 6 to 10). These arrested cells had condensed chromosomes with dispersed Golgi (Figure 5D, NR). STING had a cytosolic but granular distribution in arrested cells, which did not change upon pGL3 transfection. In contrast, STING clearly localized to p230-positive Golgi structures upon pGL3 transfection of G_1_ cells (Figure 5D, E). On average, across four independent synchronization experiments, pGL3-dependent phosphorylation of IRF3 was attenuated during mitosis, only becoming robust at 180 min (Figure 5C), when the bulk population of cells reached G_1_ and had intact Golgi with clear STING localization upon pGL3 transfection (Figure 5D, E).

### cGAS and STING are Non-Responsive During Mitosis

Recent work has shown that upon NEBD, cGAS localizes to mitotic chromosomes via direct binding to H2a/H2b dimers [53]. However this binding is not via the DNA-binding domains of cGAS that underlie DNA-dependent activation, resulting in a relatively low production of cGAMP. Thus, chromatin appears to blunt cGAS activation [34, 53], suggesting a means for the cell to avoid cGAS-driven IFN responses to self chromosomes during open mitosis. One unexplored aspect is whether chromatin-bound cGAS, or the pool of cGAS that might remain cytosolic during open mitosis, would still be responsive to foreign (naked) DNA–and whether that activation would then cause IRF3 phosphorylation.

We observed only low activation of IRF3 in response to pGL3 transfection in NOC-arrested cells (Figure 5B,C and 6A). To determine if the dampening of the pathway in these arrested cells was at the level of cGAS or STING, we measured cGAMP production in response to pGL3 transfection and examined cGAS subcellular distribution. Similar to pIRF3, cGAMP production was blunted in transfected arrested cells (NR) compared to asynchronous interphase cells (Figure 6B). Confocal microscopy of asynchronous cells revealed that cGAS was mostly nuclear although some signal was evident within the cytosol (Figure 6C), in agreement with a recent report [54]. Within arrested cells, the vast majority of cGAS was chromatin-bound and Golgi were well-dispersed (Figure 6D). These results agree with recent work showing that cGAS is predominantely nuclear, regardless of cell cycle phase or activation status [54]. Combined, our data suggest that chromatin-bound cGAS is unable to produce a robust cGAMP response to either chromosomes or transfected DNA.

**Figure 6.**
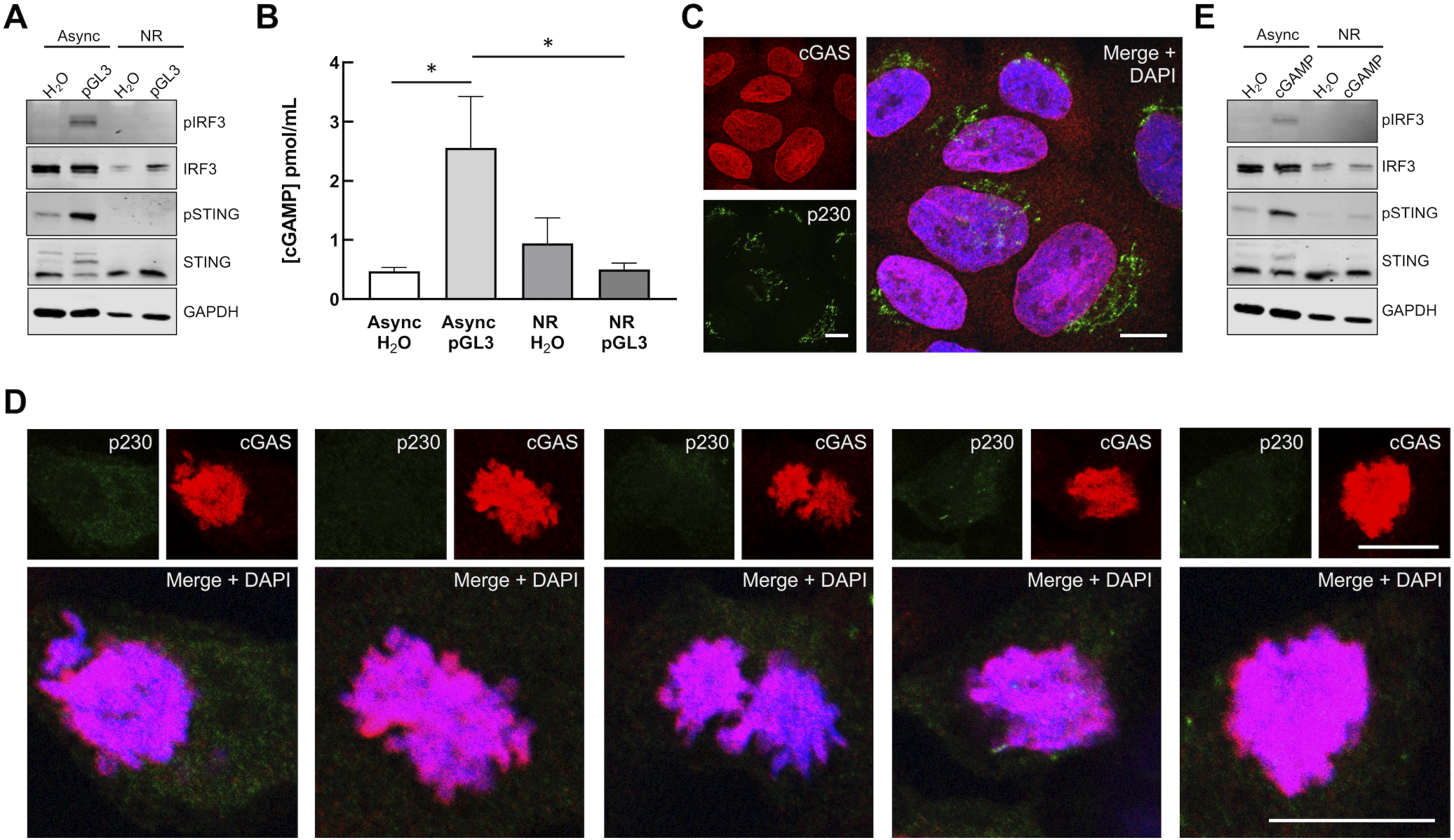
cGAS and STING are non-responsive during mitosis. **(A)** Mitotic phospho-IRF3 and phospho-STING responses to pGL3. Asynchronous and arrested cells were stimulated by transfection of 500 ng pGL3 or water for 90 min prior to western blots. **(B)** Mitotic cGAMP responses to pGL3. Asynchronous and arrested cells were transfected with 500 ng pGL3. Non-internalized transfection complexes were removed by media change 2 hr after the initial transfection. Five hours post-transfection, cGAMP levels were measured by ELISA. *p < 0.05, n = 2 biological replicates, with n = 2 technical replicates each. **(C, D)** cGAS subcellular localization and Golgi morphology in asynchronous **(C)** and arrested **(D)** HaCaT cells. Scale bars = 10 μm. **(E)** Mitotic phospho-IRF3 and phospho-STING responses to exogenous cGAMP. Asynchronous and arrested cells were treated with 25 μg/mL 2’-3’cGAMP for 2 hr prior to western blots.

Although we detected slightly elevated levels of cGAMP in unstimulated arrested versus asynchronous cells (Figure 6B), the subtle difference was not significant, consistent with recent work showing cGAS generates only low levels of cGAMP upon activation by chromatin [34, 53]. Mitotic cells could potentially take up exogenous cGAMP via SLC19A1 [84, 85] or LRRC8 [86] transporters, or directly from neighboring cells [87, 88]. Recent work has revealed a primordial role for cGAMP-dependent STING activation in triggering autophagy through WIPI2/ATG5 in a TBK1/IRF3-independent manner [75]. These Golgi-derived autophagosomes promote clearance of cytosolic DNA and incoming DNA viruses like HSV1 [75]. Although mitotic cells appear to avoid classical induction of autophagy via CDK1 phosphorylation of ATG13, ULK1, and ATG14 [89, 90], downstream STING-dependent activation of autophagophore formation/elongation via WIPI2/ATG5 during open mitosis could be deleterious to daughter cell survival [91, 92].

Considering the potential for activation of STING by low levels of endogenous cGAMP or uptake of exogenous cGAMP, we assessed whether mitotic Golgi vesiculation would prevent STING activation by exogenous cGAMP. Neither transfection of pGL3 nor addition of exogenous cGAMP stimulated pIRF3 or pSTING in arrested cells (Figure 6E), suggesting that Golgi vesiculation reinforces cGAS inactivation as a secondary barrier to cGAS/STING activation during open mitosis. During revision of this manuscript a paper was published describing how cGAS is phosphorylated and inactivated by the kinase Cdk1-cyclin B during mitosis [93]. Using different cell types their findings largely agree with what we observe herein-that chromosome-bound cGAS is inactive to stimulation by exogenous transfected DNA in mitosis and that STING is non-responsive to exogenous cGAMP in mitotic cells. Interestingly, when mitotic cGAS inactivation was blunted by use of the Cdk1 inhibitor RO-3306, despite increased cGAMP levels or addition of exogenous cGAMP, there was still lack of activated pSTING and pIRF3 [93], consistent with our data suggesting pathway inhibition by a vesiculated Golgi. Fragmentation and vesiculation of the Golgi is a natural G_2_/M checkpoint for mitotic progression [55, 94–96], thus we are unable to experimentally prevent Golgi dispersal during mitosis to determine if cGAMP-dependent STING activity would then be restored.

Given that cGAS/STING-dependent elevation of pIRF3 during mitosis (particularly during prolonged mitosis) can induce apoptosis [53] and activated STING can induce a potentially harmful autophagic response [75], a parallel dampening mechanism like Golgi dispersal likely serves an important role to limit potentially harmful cGAS/STING signaling during open mitosis. Further, many other viruses and bacteria induce dramatic alteration of Golgi membranes during infection [97–101]. Microbial alteration of Golgi integrity may be an important unrecognized tactic to blunt host cGAS/STING responses.

## Materials & Methods

### Tissue Culture

HaCaT cells were grown in high glucose DMEM (Gibco 11965-092) supplemented with 10% FBS (Gibco A31606-02) and antibiotic-antimycotic (ThermoFisher 15240062). Cells were cultured at 37°C with 5% CO_2_ and passaged every 2-3 days to maintain subconfluence.

### Nucleic Acid Transfections

HaCaTs were plated at 60,000 cells per well in a 24-well plate. Cells were transfected with 500 ng dsDNA oligonucleotide HSV-60 (InvivoGen tlrl-hsv60n, 60bp), calf thymus DNA (Sigma D4764, >20 kb), or endotoxin-free pGL3 (Promega E1751, 5.3 kb), or 500 ng high molecular weight poly(I:C) (InvivoGen tlrl-pic, 1.5 kb to 8 kb) using Lipofectamine 2000 (ThermoFisher 11668) in OptiMEM (Life Technologies). At various timepoints post-transfection, cells were washed once with PBS and lysed in 1x RIPA lysis buffer (50 mM Tris-HCl pH 8.0, 150 mM NaCl, 1% NP40, 0.5% sodium deoxycholate, 0.1% SDS), supplemented with 1x reducing SDS-PAGE loading buffer (62.5 mM Tris pH 6.8, 10% glycerol, 2% SDS, 0.5% bromophenol blue, 5% β-mercaptoethanol), 1x protease inhibitor cocktail (Sigma P1860), 1 mM PMSF, and 1x PhosSTOP phosphatase inhibitor cocktail (Roche 04906845001). Samples were boiled for 5 min at 95°C and stored at −80°C until separated and analyzed by SDS-PAGE.

### siRNA Experiments

Pooled scramble (sc-37007), cGAS (sc-95512), STING (sc-92042), and IRF3 (sc-35710) siRNA duplexes were obtained from Santa Cruz Biotechnologies. HaCaTs were plated at 30,000 cells per well in a 24-well plate with Ab/Am-free DMEM/10% FBS. The following day, cells were washed twice with PBS and the PBS replaced with OptiMEM. Cells were transfected with 50 nM siRNA using Lipofectamine RNAiMax (Life Technologies 13778150). At 16 hr post-siRNA transfection, cells were washed twice with PBS and the PBS was replaced with Ab/Am-free DMEM/10% FBS. Cells were transfected with pGL3 24 hr post-siRNA transfection, as described above. At 90 min post-transfection, samples were collected for western blotting as described above.

### Golgi Disruption

HaCaTs were plated at 60,000 cells per well on glass coverslips in 24-well plates. The following day, cells were treated with 5 μM nocodazole (Santa Cruz sc-3518), 10 μM golgicide A (Santa Cruz sc-215103), or 150 nM brefeldin A (Sigma B6542), or DMSO vehicle for 1 hr prior to further experimental treatments, and drugs were left on for the duration of these experiments.

### Cell Synchronization

HaCaTs were plated at 7.5 million cells on a 10 cm dish in 10% FBS/DMEM and allowed to reach 100% confluence. The day after plating, the media was changed to 1% FBS/DMEM and cells remained under contact inhibition at low-serum conditions for 48 hr to synchronize at G_0_. Cells were then replated at 30,000 cells per well on 24-well plates with or without glass cloverslips in 10% FBS/DMEM and allowed to progress through S phase. After 24 hr, cells were treated with 50 ng/mL nocodazole (Santa Cruz sc-3518) for 12 hr to synchronize at prometaphase, at which point they were released with two PBS washes and incubated in 10% FBS/DMEM. Cells were transfected with pGL3 as described above, and either harvested for western blotting or prepared for immunofluorescence or cell cycle analysis as described.

### Cell Cycle Analysis

Cell cycle status was analyzed by propidium iodide (PI) incorporation and flow cytometry. HaCaTs synchronized in the manner described above. At various time points during the synchronization and release from prometaphase, cells were collected by trypsinization and pelleted at 1000 rpm for 10 min at 4°C. The pellet was resuspended in ice cold 70% ethanol to fix the cells and stored at −20°C until ready for PI staining. Cells were pelleted at 2000 rpm for 15 min at 4°C, resuspended in PBS, pH 7.4 containing 40 μg/mL PI and 500 μg/mL RNase A, and incubated at 37°C for 30 min. PI-stained cells were analyzed using the BD Biosciences FACSCanto-II flow cytometer and Diva 8.0 software.

### Immunofluorescence

HaCaTs were plated at 60,000 cells per well on glass coverslips in 24-well plates. The following day, cells were treated treated and transfected as described above. For mitotic sync experiments: HaCaTs were syncronized to pro-metaphase as described above. For all experiments, following transfection, cells were fixed with 2% paraformaldehyde/PBS for 10 min at room temperature (RT) and permeabilized with 0.2% Triton X-100/PBS for 10 min at RT. Samples were blocked in 4% BSA/1% goat serum/PBS overnight at 4°C. Rabbit polyclonal anti-TGN46 (Sigma T7576. 1:500), mouse monoclonal anti-p230 (BD Biosciences 611280, 1:500), mouse monoclonal anti-IRF3 (Abcam ab50772, 1:100), rabbit anti-cGAS (Cell Signaling 15102, 1:100), and rabbit monoclonal anti-STING (Abcam ab181125, 1:500) were used as primary antibodies. Alexa Fluor-488, -555 and -647 labeled goat anti-mouse and goat anti-rabbit secondary antibodies (Life Technologies A11029, A21424, A21429, and A21236) were used at 1:1,000. Samples were then stained with 4′,6-diamidino-2-phenylindole (DAPI) (Sigma Aldrich D9542-10MG) at 1μg/mL for 30 sec. Coverslips were mounted on glass slides with Prolong Antifade Diamond (Life Technologies P36970) and analyzed by confocal microscopy.

### Confocal Microscopy

Following the preparation of immunofluorescence slides, confocal microscopy was performed using a Zeiss LSM880 system with 405 nm, 488 nm, and 543 nm lasers. Samples were examined using an oiled 63x objective, and Z-stacks with a 0.32 μm depth per plane were taken of each image. Representative single-plane images and Z-stacks were processed with the Zen Blue software.

### Colocalization Analysis

Manders’ overlap coefficients [102] for a STING:p230 and IRF3:DAPI channels within individual Z-stacks were determined using the JACoP plugin [103] on ImageJ [104]. Manual thresholds were set below saturation. Individual Manders overlap coefficient values and mean values from multiple Z-stacks (each containing multiple cells), across multiple fields of view, were plotted with GraphPad Prism software.

### cGAMP Stimulation

HaCaTs were plated at 60,000 asynchronous cells or 130,000 synchronous cells per well in 24-well plates. Cells were treated with 12.5 or 25 μg/mL 2’-3’cGAMP (Invivogen tlrl-nacga23) for 2 hr as indicated, then harvested for western blotting as described above.

### SDS-PAGE & Western Blotting

Samples were resolved by SDS-PAGE and transferred onto a 0.45 μm nitrocellulose membrane. Rabbit monoclonal anti-GAPDH (Cell Signaling 2118, 1:5000), mouse monoclonal anti-IRF3 (Abcam 50772, 1:100), rabbit monoclonal anti-TBK1 (Cell Signaling 3504, 1:1000), and rabbit monoclonal anti-STING (Cell Signaling 13647, 1:1000) blots were blocked in 5% non-fat powdered milk dissolved in Tris-buffered saline containing 0.1% Tween (TBST). Rabbit monoclonal anti-phospho-IRF3 (Ser396, Cell Signaling 4947, 1:1000), rabbit monoclonal anti-phospho-STING (Ser366, Cell Signaling 19781, 1:1000), rabbit monoclonal anti-phospho-TBK1 (Ser172, Cell Signaling 5483, 1:1000), and rabbit monoclonal anti-phospho-H3 (Ser10, Cell Signaling 3377, 1:10,000) blots were blocked in 100% Odyssey blocking buffer (Licor 927–40000). Goat anti-rabbit DyLight 680 (Pierce 35568), goat anti-mouse DyLight 680 (Pierce 35518), goat anti-rabbit DyLight 800 (Pierce 535571) and goat anti-mouse DyLight 800 (Pierce 35521) were used as secondary antibodies at 1:10,000 in either 50% Odyssey blocking buffer/TBST or 5% milk/TBST. Blots were imaged with the Licor Odyssey Infrared Imaging System. Band intensities were quantified by densitometry using ImageJ v1.52a [104].

### cGAMP ELISA

Following Golgi vesiculation and mitotic sync experiments, cells were washed 1x with PBS and prepared for cGAMP quantification by the 2′,3′-Cyclic GAMP Direct ELISA Kit (Arbor Assays K067-H1), following the manufacturer’s protocol. cGAMP concentrations were normalized by total protein concentration, as determined by BCA Assay (Thermofisher 23225).

### RT-qPCR experiments

Following Golgi disruption with GCA and transfection with pGL3, total RNA was prepared from HaCaTs using the Qiagen RNeasy Mini Kit (Qiagen 74104). RNA was isolated at 4 hr and 8 hr post-pGL3 transfection. In the final step, RNA was eluted in 45 μL RNase/DNase free water. RNA samples were purified using the TURBO DNA-free Kit (Life Technologies AM1907), and cDNA was prepared using the High Capacity cDNA Reverse Transcription Kit (ThermoFisher Scientific 4368814). To prepare cDNA, 1 μg of RNA was used per 40 μL final reaction volume, yielding an estimated cDNA concentration of 25 ng/μL. cDNA was diluted to 3.3 ng/μL with RNase/DNase free water prior to use in qPCR, Reactions with specific ISG/IFN or TATA-binding protein (TBP) housekeeper primers were prepared using the PowerUp SYBR Green Master Mix kit (ThermoFisher Scientific A25742) and loaded onto a 384-well plate. The reactions were run on a QuantStudio6 Flex Real-Time PCR System (ThermoFisher Scientific). Delta-delta-cycle threshold (ΔΔCt) was determined relative to vehicle treated samples. Viral RNA levels were normalized to TBP housekeeper and depicted as fold change over vehicle treated samples. Error bars indicate the SEM from n = 3 technical replicates. Primer sequences are as follows: TBP-for; 5’-TAAACTTGACCTAAAGACCATTGCA-3’, TBP-rev; 5’-CAGCAAACCGCTTGGGATTA-3’, IFNB1-for; 5’-CTTGGATTCCTACAAAGAAGCAGC-3’, IFNB1-rev; 5’-TCCTCCTTCTGGAACTGCTGCA-3’, viperin-for; 5’-CCAGTGCAACTACAAATGCGGC-3’, viperin-rev, 5’-CGGTCTTGAAGAAATGGCTCTCC-3’, IFI6-for; 5’-TGATGAGCTGGTCTGCGATCCT-3’, IFI6-rev; 5’-GTAGCCCATCAGGGCACCAATA-3’, HERC5- for; 5’-CAACTGGGAGAGCCTTGTGGTT-3’, HERC5-rev; 5’-CTGGACCAGTTTGCTGAAAGTGG-3’, IFIT2-for; 5’-GGAGCAGATTCTGAGGCTTTGC-3’, IFIT2-rev; 5’-GGATGAGGCTTCCAGACTCCAA-3’, IFIT3-for; 5’-CCTGGAATGCTTACGGCAAGCT-3’, IFIT3-rev; 5’-GAGCATCTGAGAGTCTGCCCAA-3’, CXCL10-for; 5’-GGTGAGAAGAGATGTCTGAATCC-3’, CXCL10-rev; 5’-GTCCATCCTTGGAAGCACTGCA-3’, CXCL11-for; 5’-AAGGACAACGATGCCTAAATCCC-3’, CXCL11-rev; 5’-CAGATGCCCTTTTCCAGGACTTC-3’.

### Statistics

Statistical analyses were performed using Prism 6 (GraphPad Software). Statistics for the pIRF3 blot densitometry from Golgi vesiculation experiments were determined by an unpaired t-test with the Holm-Sidak correction. Statistics for the cGAMP ELISAs and pIRF3 densitometry in Figure 5 were determined by ordinary one-way ANOVA with the Tukey correction for multiple comparisons. RT-qPCR data were analyzed by two-way ANOVA with Tukey’s multiple comparison. Colocalization statistics were calculated using a two-sample unpaired t-test as recommended for colocalization analysis [105]. Error bars on graphs represent standard error of mean. Number of biological and technical replicates are noted in figure legends.

## Acknowledgements

We are grateful to Jim DeCaprio for advice on cell synchronization, Koenraad Van Doorslaer, Robert Jackson, and David Williams for their assistance with RT-qPCR, helpful discussion, and critical reading of this manuscript, and Joshua Uhlorn for discussion and advice on statistics. We thank Anne Cress for the HaCaT cell line. We thank the UA BIO5 Media Facility, Patty Jansma of the UA ORD Imaging Core-Marley, and John Fitch of the UACC/ARL Cytometry Core Facility. This work was supported by grant R01-AI10875 from the National Institute for Allergy and Infectious Diseases, grant 1R01GM136853 from the National Institute for General Medical Sciences, and by a grant from the Sloan Scholars Mentoring Network of the Social Science Research Council with funds provided by the Alfred P. Sloan Foundation. B.L.U. is a graduate student supported by the National Science Foundation Graduate Research Fellowship Grant DGE-1143953.

## References

1. Medzhitov, R., Recognition of microorganisms and activation of the immune response. Nature, 2007. 449(7164): p. 819–26.

2. Takeuchi, O. and S. Akira, Pattern recognition receptors and inflammation. Cell, 2010. 140(6): p. 805–20.

3. Stetson, D.B. and R. Medzhitov, Type I interferons in host defense. Immunity, 2006. 25(3): p. 373–81.

4. Wu, J. and Z.J. Chen, Innate immune sensing and signaling of cytosolic nucleic acids. Annu Rev Immunol, 2014. 32: p. 461–88.

5. Li, T. and Z.J. Chen, The cGAS-cGAMP-STING pathway connects DNA damage to inflammation, senescence, and cancer. J Exp Med, 2018. 215(5): p. 1287–1299.

6. Chen, Q., L. Sun, and Z.J. Chen, Regulation and function of the cGAS-STING pathway of cytosolic DNA sensing. Nat Immunol, 2016. 17(10): p. 1142–9.

7. Tao, J., X. Zhou, and Z. Jiang, cGAS-cGAMP-STING: The three musketeers of cytosolic DNA sensing and signaling. IUBMB Life, 2016. 68(11): p. 858–870.

8. Stetson, D.B. and R. Medzhitov, Recognition of cytosolic DNA activates an IRF3-dependent innate immune response. Immunity, 2006. 24(1): p. 93–103.

9. Sun, L., et al., Cyclic GMP-AMP synthase is a cytosolic DNA sensor that activates the type I interferon pathway. Science, 2013. 339(6121): p. 786–91.

10. Ishikawa, H. and G.N. Barber, STING is an endoplasmic reticulum adaptor that facilitates innate immune signalling. Nature, 2008. 455(7213): p. 674–8.

11. Ishikawa, H., Z. Ma, and G.N. Barber, STING regulates intracellular DNA-mediated, type I interferon-dependent innate immunity. Nature, 2009. 461(7265): p. 788–92.

12. Yin, Q., et al., Cyclic di-GMP sensing via the innate immune signaling protein STING. Mol Cell, 2012. 46(6): p. 735–45.

13. Gao, P., et al., Structure-function analysis of STING activation by c[G(2’,5’)pA(3’,5’)p] and targeting by antiviral DMXAA. Cell, 2013. 154(4): p. 748–62.

14. Ogawa, E., et al., The binding of TBK1 to STING requires exocytic membrane traffic from the ER. Biochem Biophys Res Commun, 2018. 503(1): p. 138–145.

15. Tanaka, Y. and Z.J. Chen, STING specifies IRF3 phosphorylation by TBK1 in the cytosolic DNA signaling pathway. Sci Signal, 2012. 5(214): p. ra20.

16. Abe, T. and G.N. Barber, Cytosolic-DNA-mediated, STING-dependent proinflammatory gene induction necessitates canonical NF-kappaB activation through TBK1. J Virol, 2014. 88(10): p. 5328–41.

17. Honda, K. and T. Taniguchi, IRFs: master regulators of signalling by Toll-like receptors and cytosolic pattern-recognition receptors. Nat Rev Immunol, 2006. 6(9): p. 644–58.

18. Luo, W.W., et al., iRhom2 is essential for innate immunity to DNA viruses by mediating trafficking and stability of the adaptor STING. Nat Immunol, 2016. 17(9): p. 1057–66.

19. Sun, M.S., et al., TMED2 Potentiates Cellular IFN Responses to DNA Viruses by Reinforcing MITA Dimerization and Facilitating Its Trafficking. Cell Rep, 2018. 25(11): p. 3086–3098 e3.

20. Srikanth, S., et al., The Ca(2+) sensor STIM1 regulates the type I interferon response by retaining the signaling adaptor STING at the endoplasmic reticulum. Nat Immunol, 2019. 20(2): p. 152–162.

21. Li, Y., et al., TMEM203 is a binding partner and regulator of STING-mediated inflammatory signaling in macrophages. Proc Natl Acad Sci U S A, 2019. 116(33): p. 16479–16488.

22. Saitoh, T., et al., Atg9a controls dsDNA-driven dynamic translocation of STING and the innate immune response. Proc Natl Acad Sci U S A, 2009. 106(49): p. 20842–6.

23. Mukai, K., et al., Activation of STING requires palmitoylation at the Golgi. Nat Commun, 2016. 7: p. 11932.

24. Tsuchida, T., et al., The ubiquitin ligase TRIM56 regulates innate immune responses to intracellular double-stranded DNA. Immunity, 2010. 33(5): p. 765–76.

25. Zhang, J., et al., TRIM32 protein modulates type I interferon induction and cellular antiviral response by targeting MITA/STING protein for K63-linked ubiquitination. J Biol Chem, 2012. 287(34): p. 28646–55.

26. Shang, G., et al., Cryo-EM structures of STING reveal its mechanism of activation by cyclic GMP-AMP. Nature, 2019. 567(7748): p. 389–393.

27. Ergun, S.L., et al., STING Polymer Structure Reveals Mechanisms for Activation, Hyperactivation, and Inhibition. Cell, 2019. 178(2): p. 290–301 e10.

28. Zhang, C., et al., Structural basis of STING binding with and phosphorylation by TBK1. Nature, 2019. 567(7748): p. 394–398.

29. Chen, W., et al., ER Adaptor SCAP Translocates and Recruits IRF3 to Perinuclear Microsome Induced by Cytosolic Microbial DNAs. PLoS Pathog, 2016. 12(2): p. e1005462.

30. Ablasser, A. and Z.J. Chen, cGAS in action: Expanding roles in immunity and inflammation. Science, 2019. 363(6431).

31. Bakhoum, S.F., et al., Chromosomal instability drives metastasis through a cytosolic DNA response. Nature, 2018. 553(7689): p. 467–472.

32. Gluck, S., et al., Innate immune sensing of cytosolic chromatin fragments through cGAS promotes senescence. Nat Cell Biol, 2017. 19(9): p. 1061–1070.

33. Harding, S.M., et al., Mitotic progression following DNA damage enables pattern recognition within micronuclei. Nature, 2017. 548(7668): p. 466–470.

34. Mackenzie, K.J., et al., cGAS surveillance of micronuclei links genome instability to innate immunity. Nature, 2017. 548(7668): p. 461–465.

35. Yang, H., et al., cGAS is essential for cellular senescence. Proc Natl Acad Sci U S A, 2017. 114(23): p. E4612–E4620.

36. Dou, Z., et al., Cytoplasmic chromatin triggers inflammation in senescence and cancer. Nature, 2017. 550(7676): p. 402–406.

37. Motwani, M., S. Pesiridis, and K.A. Fitzgerald, DNA sensing by the cGAS-STING pathway in health and disease. Nat Rev Genet, 2019. 20(11): p. 657–674.

38. Konno, H., et al., Pro-inflammation Associated with a Gain-of-Function Mutation (R284S) in the Innate Immune Sensor STING. Cell Rep, 2018. 23(4): p. 1112–1123.

39. Liu, Y., et al., Activated STING in a vascular and pulmonary syndrome. N Engl J Med, 2014. 371(6): p. 507–518.

40. Picard, C., et al., Severe Pulmonary Fibrosis as the First Manifestation of Interferonopathy (TMEM173 Mutation). Chest, 2016. 150(3): p. e65–71.

41. Saldanha, R.G., et al., A Mutation Outside the Dimerization Domain Causing Atypical STING-Associated Vasculopathy With Onset in Infancy. Front Immunol, 2018. 9: p. 1535.

42. Davidson, S., et al., An Update on Autoinflammatory Diseases: Interferonopathies. Curr Rheumatol Rep, 2018. 20(7): p. 38.

43. Stetson, D.B., et al., Trex1 prevents cell-intrinsic initiation of autoimmunity. Cell, 2008. 134(4): p. 587–98.

44. Gray, E.E., et al., Cutting Edge: cGAS Is Required for Lethal Autoimmune Disease in the Trex1-Deficient Mouse Model of Aicardi-Goutieres Syndrome. J Immunol, 2015. 195(5): p. 1939–43.

45. Yoshida, H., et al., Lethal anemia caused by interferon-beta produced in mouse embryos carrying undigested DNA. Nat Immunol, 2005. 6(1): p. 49–56.

46. Sun, C., et al., Cellular Requirements for Sensing and Elimination of Incoming HSV-1 DNA and Capsids. J Interferon Cytokine Res, 2019. 39(4): p. 191–204.

47. Hare, D.N., et al., Membrane Perturbation-Associated Ca2+ Signaling and Incoming Genome Sensing Are Required for the Host Response to Low-Level Enveloped Virus Particle Entry. J Virol, 2015. 90(6): p. 3018–27.

48. Lam, E., S. Stein, and E. Falck-Pedersen, Adenovirus detection by the cGAS/STING/TBK1 DNA sensing cascade. J Virol, 2014. 88(2): p. 974–81.

49. Wang, F., et al., S6K-STING interaction regulates cytosolic DNA-mediated activation of the transcription factor IRF3. Nat Immunol, 2016. 17(5): p. 514–522.

50. Watkinson, R.E., et al., TRIM21 Promotes cGAS and RIG-I Sensing of Viral Genomes during Infection by Antibody-Opsonized Virus. PLoS Pathog, 2015. 11(10): p. e1005253.

51. Dai, P., et al., Modified vaccinia virus Ankara triggers type I IFN production in murine conventional dendritic cells via a cGAS/STING-mediated cytosolic DNA-sensing pathway. PLoS Pathog, 2014. 10(4): p. e1003989.

52. Garcia-Belmonte, R., et al., African Swine Fever Virus Armenia/07 Virulent Strain Controls Interferon Beta Production through the cGAS-STING Pathway. J Virol, 2019. 93(12).

53. Zierhut, C., et al., The Cytoplasmic DNA Sensor cGAS Promotes Mitotic Cell Death. Cell, 2019. 178(2): p. 302–315 e23.

54. Volkman, H.E., et al., Tight nuclear tethering of cGAS is essential for preventing autoreactivity. Elife, 2019. 8.

55. Sutterlin, C., et al., Fragmentation and dispersal of the pericentriolar Golgi complex is required for entry into mitosis in mammalian cells. Cell, 2002. 109(3): p. 359–69.

56. Tang, D. and Y. Wang, Cell cycle regulation of Golgi membrane dynamics. Trends Cell Biol, 2013. 23(6): p. 296–304.

57. Boukamp, P., et al., Normal keratinization in a spontaneously immortalized aneuploid human keratinocyte cell line. J Cell Biol, 1988. 106(3): p. 761–71.

58. Bernard, E., et al., Human keratinocytes restrict chikungunya virus replication at a post-fusion step. Virology, 2015. 476: p. 1–10.

59. Griffin, L.M., et al., Human keratinocyte cultures in the investigation of early steps of human papillomavirus infection. Methods Mol Biol, 2014. 1195: p. 219–38.

60. Kim, J.A., et al., Insights into ZIKV-Mediated Innate Immune Responses in Human Dermal Fibroblasts and Epidermal Keratinocytes. J Invest Dermatol, 2019. 139(2): p. 391–399.

61. Lim, P.Y., et al., Keratinocytes are cell targets of West Nile virus in vivo. J Virol, 2011. 85(10): p. 5197–201.

62. Lopez-Gonzalez, M., et al., Human keratinocyte cultures (HaCaT) can be infected by DENV, triggering innate immune responses that include IFNlambda and LL37. Immunobiology, 2018. 223(11): p. 608–617.

63. Puiprom, O., et al., Characterization of chikungunya virus infection of a human keratinocyte cell line: role of mosquito salivary gland protein in suppressing the host immune response. Infect Genet Evol, 2013. 17: p. 210–5.

64. Rahn, E., et al., Entry pathways of herpes simplex virus type 1 into human keratinocytes are dynamin- and cholesterol-dependent. PLoS One, 2011. 6(10): p. e25464.

65. Kisich, K.O., et al., The constitutive capacity of human keratinocytes to kill Staphylococcus aureus is dependent on beta-defensin 3. J Invest Dermatol, 2007. 127(10): p. 2368–80.

66. Petrucelli, M.F., et al., Dual RNA-Seq Analysis of Trichophyton rubrum and HaCat Keratinocyte Co-Culture Highlights Important Genes for Fungal-Host Interaction. Genes (Basel), 2018. 9(7).

67. Almine, J.F., et al., IFI16 and cGAS cooperate in the activation of STING during DNA sensing in human keratinocytes. Nat Commun, 2017. 8: p. 14392.

68. Dunphy, G., et al., Non-canonical Activation of the DNA Sensing Adaptor STING by ATM and IFI16 Mediates NF-kappaB Signaling after Nuclear DNA Damage. Mol Cell, 2018. 71(5): p. 745–760 e5.

69. Kim, J.A., et al., STING Is Involved in Antiviral Immune Response against VZV Infection via the Induction of Type I and III IFN Pathways. J Invest Dermatol, 2017. 137(10): p. 2101–2109.

70. Olagnier, D., et al., Nrf2 negatively regulates STING indicating a link between antiviral sensing and metabolic reprogramming. Nat Commun, 2018. 9(1): p. 3506.

71. Skouboe, M.K., et al., STING agonists enable antiviral cross-talk between human cells and confer protection against genital herpes in mice. PLoS Pathog, 2018. 14(4): p. e1006976.

72. Cole, N.B., et al., Golgi dispersal during microtubule disruption: regeneration of Golgi stacks at peripheral endoplasmic reticulum exit sites. Mol Biol Cell, 1996. 7(4): p. 631–50.

73. Helms, J.B. and J.E. Rothman, Inhibition by brefeldin A of a Golgi membrane enzyme that catalyses exchange of guanine nucleotide bound to ARF. Nature, 1992. 360(6402): p. 352–4.

74. Saenz, J.B., et al., Golgicide A reveals essential roles for GBF1 in Golgi assembly and function. Nat Chem Biol, 2009. 5(3): p. 157–65.

75. Gui, X., et al., Autophagy induction via STING trafficking is a primordial function of the cGAS pathway. Nature, 2019. 567(7747): p. 262–266.

76. Kawai, T., et al., IPS-1, an adaptor triggering RIG-I- and Mda5-mediated type I interferon induction. Nat Immunol, 2005. 6(10): p. 981–8.

77. Yoneyama, M., et al., The RNA helicase RIG-I has an essential function in double-stranded RNA-induced innate antiviral responses. Nat Immunol, 2004. 5(7): p. 730–7.

78. Farmaki, T., et al., Forward and retrograde trafficking in mitotic animal cells. ER-Golgi transport arrest restricts protein export from the ER into COPII-coated structures. J Cell Sci, 1999. 112 (Pt 5): p. 589–600.

79. Kreiner, T. and H.P. Moore, Membrane traffic between secretory compartments is differentially affected during mitosis. Cell Regul, 1990. 1(5): p. 415–24.

80. Prescott, A.R., et al., Evidence for prebudding arrest of ER export in animal cell mitosis and its role in generating Golgi partitioning intermediates. Traffic, 2001. 2(5): p. 321–35.

81. Warren, G., et al., Newly synthesized G protein of vesicular stomatitis virus is not transported to the cell surface during mitosis. J Cell Biol, 1983. 97(5 Pt 1): p. 1623–8.

82. Hughes, H. and D.J. Stephens, Sec16A defines the site for vesicle budding from the endoplasmic reticulum on exit from mitosis. J Cell Sci, 2010. 123(Pt 23): p. 4032–8.

83. Ward, T.H., et al., Maintenance of Golgi structure and function depends on the integrity of ER export. J Cell Biol, 2001. 155(4): p. 557–70.

84. Luteijn, R.D., et al., SLC19A1 transports immunoreactive cyclic dinucleotides. Nature, 2019. 573(7774): p. 434–438.

85. Ritchie, C., et al., SLC19A1 Is an Importer of the Immunotransmitter cGAMP. Mol Cell, 2019. 75(2): p. 372–381 e5.

86. Zhou, C., et al., Transfer of cGAMP into Bystander Cells via LRRC8 Volume-Regulated Anion Channels Augments STING-Mediated Interferon Responses and Anti-viral Immunity. Immunity, 2020. 52(5): p. 767–781 e6.

87. Ablasser, A., et al., Cell intrinsic immunity spreads to bystander cells via the intercellular transfer of cGAMP. Nature, 2013. 503(7477): p. 530–4.

88. Fykerud, T.A., et al., Mitotic cells form actin-based bridges with adjacent cells to provide intercellular communication during rounding. Cell Cycle, 2016. 15(21): p. 2943–2957.

89. Odle, R.I., et al., An mTORC1-to-CDK1 Switch Maintains Autophagy Suppression during Mitosis. Mol Cell, 2019.

90. Willson, J., Mitosis flips the switch on autophagy control. Nat Rev Mol Cell Biol, 2019.

91. Eskelinen, E.L., et al., Inhibition of autophagy in mitotic animal cells. Traffic, 2002. 3(12): p. 878–93.

92. Mathiassen, S.G., D. De Zio, and F. Cecconi, Autophagy and the Cell Cycle: A Complex Landscape. Front Oncol, 2017. 7: p. 51.

93. Zhong, L., et al., Phosphorylation of cGAS by CDK1 impairs self-DNA sensing in mitosis. Cell Discov, 2020. 6: p. 26.

94. Cervigni, R.I., et al., JNK2 controls fragmentation of the Golgi complex and the G2/M transition through phosphorylation of GRASP65. J Cell Sci, 2015. 128(12): p. 2249–60.

95. Colanzi, A., et al., The Golgi mitotic checkpoint is controlled by BARS-dependent fission of the Golgi ribbon into separate stacks in G2. EMBO J, 2007. 26(10): p. 2465–76.

96. Corda, D., et al., Golgi complex fragmentation in G2/M transition: An organelle-based cell-cycle checkpoint. IUBMB Life, 2012. 64(8): p. 661–70.

97. Canton, J. and P.E. Kima, Interactions of pathogen-containing compartments with the secretory pathway. Cell Microbiol, 2012. 14(11): p. 1676–86.

98. Machamer, C.E., The Golgi complex in stress and death. Front Neurosci, 2015. 9: p. 421.

99. Mousnier, A., et al., Human rhinovirus 16 causes Golgi apparatus fragmentation without blocking protein secretion. J Virol, 2014. 88(20): p. 11671–85.

100. Heuer, D., et al., Chlamydia causes fragmentation of the Golgi compartment to ensure reproduction. Nature, 2009. 457(7230): p. 731–5.

101. Pierini, R., et al., Modulation of membrane traffic between endoplasmic reticulum, ERGIC and Golgi to generate compartments for the replication of bacteria and viruses. Semin Cell Dev Biol, 2009. 20(7): p. 828–33.

102. Manders, E.M.M., F.J. Verbeek, and J.A. Aten, Measurement of co-localization of objects in dual-colour confocal images. Journal of Microscopy, 1993. 169(3): p. 375–382.

103. Bolte, S. and F.P. Cordelieres, A guided tour into subcellular colocalization analysis in light microscopy. J Microsc, 2006. 224(Pt 3): p. 213–32.

104. Schneider, C.A., W.S. Rasband, and K.W. Eliceiri, NIH Image to ImageJ: 25 years of image analysis. Nat Methods, 2012. 9(7): p. 671–5.

105. McDonald, J.H. and K.W. Dunn, Statistical tests for measures of colocalization in biological microscopy. J Microsc, 2013. 252(3): p. 295–302.

